# Insights into Ice-Growth Inhibition and Hydration Ordering by the Ca^2+^-Dependent Antifreeze Protein MaIBP_RIV

**DOI:** 10.64898/2026.11.24.768877

**Authors:** Sourav Satapathi, Biman Jana

## Abstract

Antifreeze proteins (AFPs) inhibit ice growth by binding to specific ice surfaces, although the molecular basis of this activity remains incompletely understood. Here, we investigate the ice-growth inhibition mechanism of the calcium-dependent antifreeze protein MaIBP_RIV from Marinomonas arctica using all-atom molecular dynamics simulations. MaIBP_RIV exhibits stable binding to the primary prism and basal ice planes through its ice-binding surface (IBS), effectively suppressing ice growth. IBS binding to the secondary prism plane is weaker and only temporarily inhibits growth. In contrast, the non-ice-binding surface (non-IBS) is repelled by the growing ice front. Our results also explain that Ca^2+^ stabilises the IBS and essential for ice-growth inhibition, while removing Ca^2+^ reduces IBS flatness, which significantly impairs its ice-growth activity. Experimentally characterised mutations on IBS residues (S44Y, G63D, N65Y, and D87Y) result in continuous ice growth, consistent with their reduced antifreeze activity. Hydration analysis of water molecules around IBS and non-IBS suggests that, by decreasing the temperature from 300K to 240K, the water molecules around IBS show more ice-like ordering compared to those around the non-IBS.

## 1. INTRODUCTION

The protection of biological samples during repeated freezing and thawing is an important challenge in cryopreservation.^1^. Natural ice-growth inhibitors and antifreeze proteins (AFPs) attract considerable attention because of their possible applications in cryopreservation, regenerative medicine, and the food industry.^2–5^. AFPs are identified in many organisms, including fish, insects, plants, bacteria, and fungi that live in cold and icy environments.^5,6^. A single organism can also produce several types of AFPs. For example, fish are reported to contain five major classes of AFPs^7^: type I, type II, type III, type IV, and antifreeze glycoproteins^8–16^. Each class can also have several different variants^17^. These AFPs show considerable structural differences and have evolved independently.

The activity of an AFP is commonly measured in terms of thermal hysteresis (TH)^18–23^. TH refers to the difference between the freezing and melting temperatures created by an AFP^6,24–26^. Insects are generally found to produce highly active AFPs, which can produce a thermal hysteresis gap of about 5–6 °C. In comparison, fish AFPs usually produce gaps of about 0.5– 1.0 °C, while plant AFPs produce gaps of approximately 0.1–0.6 °C^27–30^. Insect AFPs are described as hyperactive antifreeze proteins because of their high activity.

Despite extensive research, the exact molecular mechanism through which AFPs prevent ice growth^31^ is not completely understood. The most widely accepted explanation is the adsorption-inhibition model^32^. According to this model, an AFP attaches to the surface of growing ice and adsorbs on it. When AFPs irreversibly adsorb onto the ice surface of a growing ice plane, they create a curved ice-water interface between two bound AFPs. When the radius of this curved surface becomes smaller than the critical radius required for ice formation, further ice growth is stopped. However, when the temperature is lowered sufficiently, ice growth starts again (Gibbs-Thomson effect)^10,33^. The exact way in which AFPs attach to ice surfaces remains a topic of considerable debate.

A microfluidic study suggests that AFPs adsorb irreversibly to ice^34^. However, some recent experimental and simulation studies suggest that antifreeze glycoproteins interact with ice reversibly^35,36^. Therefore, the detailed molecular mechanism of AFP binding and the driving force involved for this adsorption remain unclear and controversial.

Several studies^37–40^ propose that interfacial water molecules at the protein-ice interface play an important role in ice recognition and binding mechanisms. Many studies suggest that the solvated water molecules around IBS play an important role in the binding and recognition phenomena^41–43^.

Meister et al. propose that long-range interactions between proteins and water contribute to the antifreeze activity of hyperactive AFPs^44^. Nutt and Smith^10^ propose another explanation known as the “preorder-binding” mechanism. They suggest that water molecules around the ice-binding surface of an AFP become more ordered. As a result, the amino acid residues present at the IBS can pre-organize the water molecules to form an ice-like structure. When the growing ice surface approaches the protein, these pre-organized water molecules reduce the energy and entropy barrier required for ice formation and promote the formation of ice between the AFP and the bulk ice. Based on X-ray crystallographic studies of AFP-ice interactions, Garnham et al. propose the “anchored clathrate water” (ACW) mechanism^45^. According to this model, AFPs organize surrounding water molecules into a structure that resembles the quasi-ice-like lattice. Hydrophobic interactions help form this quasi-ice-like arrangement^46–48^, while hydrogen bonds between the protein and water stabilize the organized water molecules. The ice-like water structure around the AFP binding site then interacts with the water layer present on the growing ice surface, eventually forming a continuous ice structure. However, some studies^41,47–49^ suggest that such pre-organization of water occurs mainly when AFPs possess large binding surfaces or form aggregates with ice-nucleating proteins (INPs). These findings challenge some aspects of the ACW and ice-like water ordering models. Other studies^50–53^ suggest that AFP molecules move relatively slowly through solution and recognize ice by matching their ice-binding sites with specific surfaces of the ice crystal. For this interaction to occur, the AFP needs to adopt the correct orientation relative to the ice surface.

Because of the practical importance of AFPs and the need to understand their basic mechanisms, researchers continue to search for new AFPs in nature. One of the most extensively studied bacterial AFPs is MpIBP^45^, which originates from Marinomonas primoryensis^48,49^, a cold-adapted bacterium isolated from Ace Lake in Antarctica. The ice-binding domain of MpIBP, known as Region IV, possesses a unique β-helical structure. Its flat ice-binding surface contains a repeated arrangement of threonine (Thr) and Asx (aspartic acid/asparagine) residues.

The ice-binding surface of MpIBP Region IV is associated with the formation of “anchored clathrate waters,” which resemble the structure of ice. This arrangement is proposed to contribute to the ability of MpIBP to recognize ice and prevent its growth. Interestingly, MpIBP is the only known AFP identified among Marinomonas species. In addition, no Marinomonas species have been found to contain DUF3494^25^, which is a common AFP domain in many bacteria. Even after extensive investigations, the researchers who discovered MpIBP did not identify any closely related AFPs.

These observations raise questions about how diverse AFPs are within the Marinomonas genus and how widely they are distributed. MaIBP shows a domain organization that is highly similar to that of MpIBP, suggesting that the two proteins might perform similar biological functions. To determine whether MaIBP has antifreeze activity and to establish whether it represents a second ice-binding protein in the Marinomonas genus, Liao et al. studied its ice-binding domain, which is referred to as MaIBP_RIV^54^. They characterized the properties and ice-binding activity of MaIBP_RIV and investigated its possible use in cell cryopreservation.

MaIBP_RIV, a calcium-dependent antifreeze protein from Marinomonas arctica, exhibits antifreeze activity through interactions between its IBS and ice surfaces. However, the molecular details governing its interaction with different ice planes and the contribution of Ca^2+^ to its antifreeze function are not fully established. Understanding these interactions is important for elucidating how AFPs selectively recognize ice and prevent its further growth.

In this study, all-atom molecular dynamics simulations are employed to investigate the ice-growth inhibition mechanism of MaIBP_RIV on the primary prism, basal and secondary prism planes of hexagonal ice. The study examines the ice growth inhibition on different planes of hexagonal ice by MaIBP_RIV, the role of Ca^2+^ in maintaining the structural integrity of the IBS, and the effects of experimentally characterized IBS mutations. In addition, hydration and ice-like ordering of water molecules around the IBS and non-ice-binding surface (non-IBS) are investigated at different temperatures. These analyses provide molecular-level insights into the contributions of specific IBS residues, Ca^2+^-dependent structural stabilization, protein-ice binding, and interfacial water organization to the antifreeze activity of MaIBP_RIV. The findings may contribute to the rational design of bioinspired antifreeze materials and cryoprotective systems.

## 2. MATERIALS AND METHODS

### 2.1 Preparation of the Systems and Simulation Details

An experimentally resolved three-dimensional structure of MaIBP_RIV from Marinomonas arctica was unavailable. The structure of MaIBP_RIV was predicted using AlphaFold 3^55^ providing the sequence of the protein. MaIBP_RIV was a calcium-dependent antifreeze protein. The Ca^2+^ ion was known to play a critical role in stabilizing the ice-binding surface (IBS) and maintaining its function. The Ca^2+^ bound MaIBP_RIV structure was subjected to energy minimization and subsequently equilibration and production molecular dynamics runs were performed. In this study, GROMACS 2020.3^56^ package was used to perform the molecular dynamics simulations using the AMBER99SB-ILDN forcefield^57,58^. For visualisation purposes, we used Visual Molecular Dynamics (VMD)^59^ package and Open Visualization Tool (OVITO)^60^. Details of the simulations are presented in Table 1.

**Table 1:** Details of the simulations performed in this study.

| System Details | Temperature(K) | ICE Plane | Box Dimension | Production Run (ns) |
|---|---|---|---|---|
| MaIBP_RIV with Ca<sup>2+</sup> (IBS)+ICE+ Water | 272K | Primary Prism Plane | 9.1*7.9*10.6 | 1000 ns |
| MaIBP_RIV with Ca<sup>2+</sup> (non-IBS) + ICE + Water | 272K | Primary Prism Plane | 9.3*8.0*10.7 | 1000 ns |
| MaIBP_RIV without Ca<sup>2+</sup> (IBS) + ICE + Water | 272K | Primary Prism Plane | 9.3*8.0*10.7 | 1000 ns |
| ICE + Water | 272K | Primary Prism Plane | 9.3*8.0*10.7 | 300 ns |
| MaIBP_RIV with Ca<sup>2+</sup> (IBS) + ICE + Water | 272K | Basal Plane | 8.5*8.7*10.7 | 1000 ns |
| MaIBP_RIV with Ca<sup>2+</sup> (non-IBS) + ICE + Water | 272K | Basal Plane | 8.5*8.7*10.7 | 1000 ns |
| MaIBP_RIV without Ca<sup>2+</sup> (IBS) + ICE + Water | 272K | Basal Plane | 8.5*8.7*10.7 | 1000 ns |
| ICE + Water | 272K | Basal Plane | 8.5*8.7*10.7 | 450 ns |
| MaIBP_RIV with Ca<sup>2+</sup> (IBS) + ICE + Water | 272K | Secondary Prism Plane | 9.3*8.0*10.7 | 1000 ns |
| MaIBP_RIV with Ca<sup>2+</sup> (non-IBS) + ICE + Water | 272K | Secondary Prism Plane | 9.3*8.0*10.7 | 1000 ns |
| MaIBP_RIV without Ca<sup>2+</sup> (IBS) + ICE + Water | 272K | Secondary Prism Plane | 9.3*8.0*10.7 | 1000 ns |
| ICE + Water | 272K | Secondary Prism Plane | 9.3*8.0*10.7 | 350 ns |
| MaIBP_RIV with Ca<sup>2+</sup> + G63D mutation (IBS) + ICE + Water | 272K | Primary Prism Plane | 9.3*8.0*10.7 | 1000 ns |
| MaIBP_RIV with Ca<sup>2+</sup> + D87Y mutation (IBS) + ICE + Water | 272K | Primary Prism Plane | 9.3*8.0*10.7 | 1000 ns |
| MaIBP_RIV with Ca<sup>2+</sup> + N65Y mutation (IBS) + ICE + Water | 272K | Primary Prism Plane | 9.3*8.0*10.7 | 1000 ns |
| MaIBP_RIV with Ca<sup>2+</sup> + S44Y mutation (IBS) + ICE + Water | 272K | Primary Prism Plane | 9.3*8.0*10.7 | 1000 ns |
| MaIBP_RIV With Ca<sup>2+</sup> + Water | 240K, 250K, 260K, 273K, 300K | No | 8.1*8.1*8.1 | 200 ns |

### 2.2. System Preparation for MaIBP_RIV AFPs

To investigate the interaction of MaIBP_RIV with different ice planes, a series of molecular dynamics simulations were performed at the ice/water interface. The simulation systems were prepared by placing the protein above an ice slab consisting of four layers of hexagonal ice (Ih). The protein is positioned such a way the distance between the ice surface and the ice-binding surface (IBS) remains 2.0 nm (Figure 1A, 1B and 1C). The IBS of MaIBP_RIV was oriented to the growing ice surface. Simulations were performed for the primary prism plane, the basal plane, and the secondary prism plane. IBS was identified as the relatively flat surface of MaIBP_RIV, consistent with previous experimental and structural studies. Unlike its Antarctic homolog MpIBP_RIV’s threonine (Thr)-rich motifs, MaIBP_RIV exhibited an irregular, Thr-free ice-binding surface (IBS). The MaIBP_RIV antifreeze protein domain contained 217 amino acid residues. It contained nine tandemly repeated GXG(N/D/S) (D/N) motifs, separated by 14-19 amino acid residues. The IBS of MaIBP_RIV contained Gly(G), Ser(S), Asn(N) and Asp(D) residues. The IBS of MaIBP_RIV is presented in the Supplementary Information (Figure S1). In another set of simulations, the non-IBS side of MaIBP_RIV (Figure S1) was oriented toward the ice surface. We also performed simulations using MaIBP_RIV structure without Ca^2+^ to explain the role of ions in IBS stability. All systems were solvated using the TIP5P^61,62^ water model. The TIP5P model was selected because it accurately reproduces several thermodynamic properties of water and exhibits a melting temperature close to the experimental melting point of ice (~273 K).

**Figure 1:**
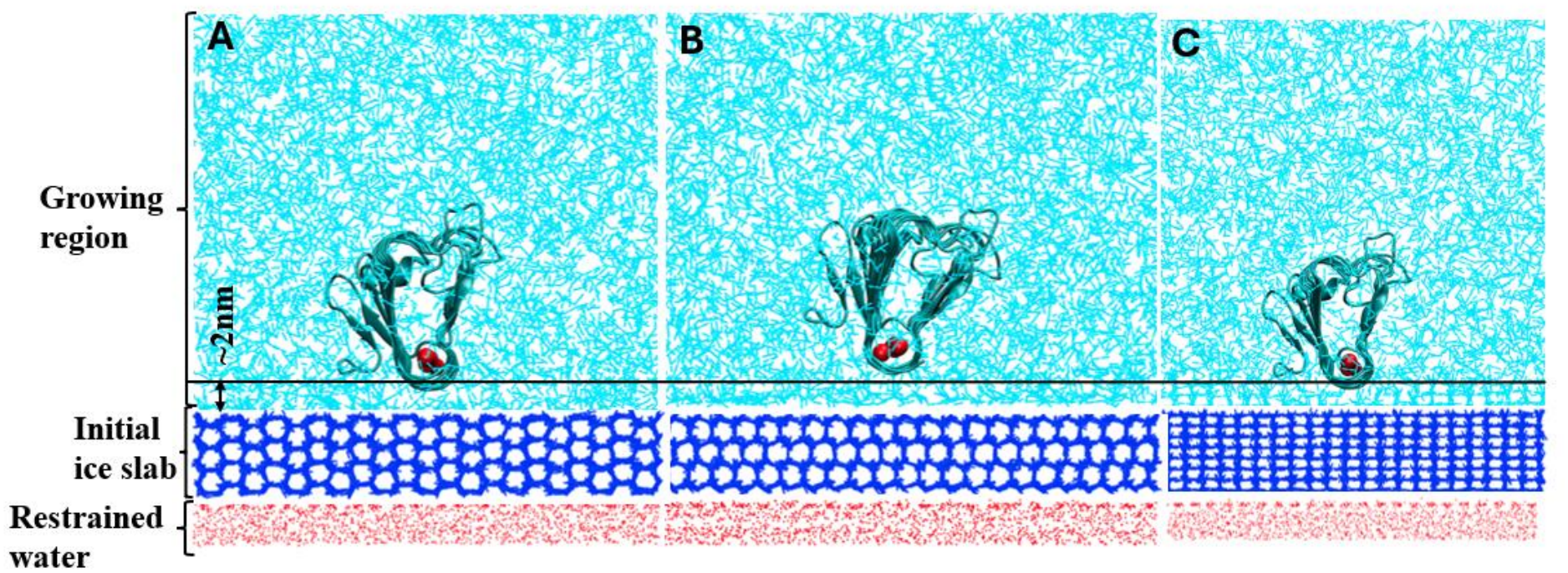
Representation of the initial structure of the MaIBP_RIV AFP placed at the ice-water interface. For the MaIBP_RIV AFP, we have considered the primary prism plane (A), basal plane (B) and secondary prism plane (C). Oxygen atoms of water molecules were shown in cyan colour. AFPs were represented in the new cartoon mode. The preformed ice surface was represented in blue colour, and the position-restrained water molecules were shown in red colour. Ca^2+^ ions were shown in VDW mode and in red colour. The ice-binding surface (IBS) of MaIBP_RIV was oriented toward the preformed hexagonal ice surface and positioned approximately 2 nm above the preformed ice slab.

### 2.3. Simulation Details of Growth Study

All systems were electrically neutralized by adding the appropriate number of counterions. Then, the systems were subjected to energy minimization using the steepest descent algorithm to remove any bad contacts and steric clashes. After energy minimization, each system was equilibrated for 1 ns in the NVT ensemble. During the equilibration phase, harmonic position restraints with a force constant of 1000 kJ mol^−1^ nm^−2^ were applied to the protein backbone, the four layers of the hexagonal ice slab, and the water molecules positioned below the ice slab. In contrast, the water molecules above the ice surface were allowed to move freely. The water molecules below the ice slab were restrained throughout the simulations to prevent ice growth in the downward direction. The final production simulations were performed in the NPT ensemble. The temperature was maintained using the Nose-Hoover^63^ thermostat with a coupling time constant of 0.5 ps. The pressure was controlled using the Parrinello-Rahman^64^ barostat with a coupling time constant of 0.5 ps. Long-range electrostatic interactions were calculated using the particle mesh Ewald (PME)^65^ method with a cutoff of 1.0 nm and a grid spacing of 0.16 nm. Van der Waals interactions were calculated using a cutoff distance of 1.0 nm. All simulations were performed with a time step of 2 fs.

### 2.4. Simulation Details for Hydration Study

The initial structure of Ca^2+^ bound MaIBP_RIV antifreeze protein (AFP) was subjected to energy minimization to remove any bad and steric contacts. Energy minimization was performed using the steepest descent algorithm. Then, the protein was solvated in the TIP5P water model. A cubic simulation box was constructed with a distance of approximately 2.0 nm between the protein surface and the box boundaries. All molecular dynamics simulations were performed using periodic boundary conditions. Short-range van der Waals interactions were calculated using a cutoff distance of 1.0 nm. Long-range electrostatic interactions were treated using the Particle Mesh Ewald (PME)^65^ method with a cutoff of 1.0 nm and a grid spacing of 0.16 nm. Next, the solvated systems were energy minimized using the steepest descent method to remove any steric clashes and bad contacts. The minimized systems were equilibrated in the NVT ensemble. The temperature was maintained using the Nose-Hoover thermostat^63^ with a coupling constant of 0.5 ps. After that, the systems were equilibrated in the NPT ensemble. During equilibration, harmonic position restraints were applied to the protein backbone atoms with a force constant of 1000 kJ mol^−1^ nm^−2^ to preserve the structure of the IBS, while water molecules and ions were allowed to move freely. The pressure was maintained using the Parrinello-Rahman barostat^64^ with a coupling constant of 0.5 ps. The final production runs were performed at temperatures of 240, 250, 260, 273, and 300 K. To investigate the temperature-dependent hydration properties around the ice-binding site (IBS), independent production simulations of 200 ns at different temperatures were performed.

Hydration analyses are performed using the final 180 ns of each trajectory, while the initial 20 ns is excluded. The structural ordering of hydration water was quantified through F_4_ order parameter. F_4_^66^ is based on the H-O-O-H torsional angle formed between hydrogen-bonded water dimers. While calculating the fraction of ordered water in terms of F_4_, we have selected the average F_4_ cutoff values of 0.2 and −0.3 for clathrate-like and ice-like water, respectively. F_4_ values of nearly 0.7 correspond to clathrate water, ~ 0.0 for liquid, and ~ −0.3 for hexagonal ice.

### 2.5. Analysis of ice growth in Simulation Trajectories

The evolution of ice growth during the simulations was calculated from the fraction of ice-like water molecules over the simulation time using the CHILL+^67^ order parameter. The CHILL+ order parameter was used to identify whether water molecules were present as liquid water, cubic ice (ice Ic), or hexagonal ice (ice Ih). The order parameter analysed the orientation of each water molecule and its four nearest neighbours. It distinguished the different phases based on the number of staggered and eclipsed bonds. In cubic ice, a water molecule forms four staggered bonds. On the other hand, in hexagonal ice, it forms three staggered and one eclipsed bond. Here we present the evolution of the fraction of ice-like molecules (cubic + hexagonal) throughout the trajectory. The order parameter has been successfully used to differentiate water ordering around protein surfaces.

## 3. RESULTS AND DISCUSSION

In the present study, we have investigated the evolution of a growing ice front in presence of MaIBP_RIV at the ice-water interface. All-atom molecular dynamics simulations were performed to provide mechanistic insights into the ice growth inhibition mechanisms by this AFP. We have selected MaIBP_RIV, a calcium-dependent antifreeze protein from Marinomonas arctica, to distinguish the effects arising from specific ice-binding surfaces. We have performed comparative simulations using the non-IBS surface of MaIBP_RIV. This surface was chosen as a control because it possesses a similar structural topology. All simulations were performed using the TIP5P water model, which was selected because it accurately reproduces several thermodynamic properties of water and exhibited a melting temperature close to the experimental melting point of ice. Ice growth simulations for MaIBP_RIV were performed at 272 K. The experimentally measured thermal hysteresis (TH) activity of MaIBP_RIV was ~1.03 °C.

### 3.1. Ice-Growth Inhibition on the Primary Prism Plane

To investigate the mechanism of ice growth inhibition by MaIBP_RIV, a series of molecular dynamics simulations were performed at the ice/water interface of different planes of hexagonal ice. Previous experimental studies suggested that MaIBP_RIV has the ability to bind multiple planes of hexagonal ice, including the prism, basal and pyramidal planes. Therefore, simulations were first performed at the primary prism plane to understand the interaction between the IBS and the growing ice surface. For comparison, simulations were also performed for the non-IBS side of MaIBP_RIV and for a pure ice/water system. In all cases (except the pure ice/water system), the protein was initially positioned approximately 20 Å above the ice surface. Representative snapshots of the simulation trajectories suggested differences in the behaviour of the three systems (Figure 2). During the initial stages of the simulation, the growing ice front propagated toward the protein surface. Within approximately 60 ns, the growing ice reached the vicinity of both the IBS and non-IBS. A similar extent of ice growth was observed in the pure ice/water system. However, significant differences were observed as the simulations progressed.

**Figure 2:**
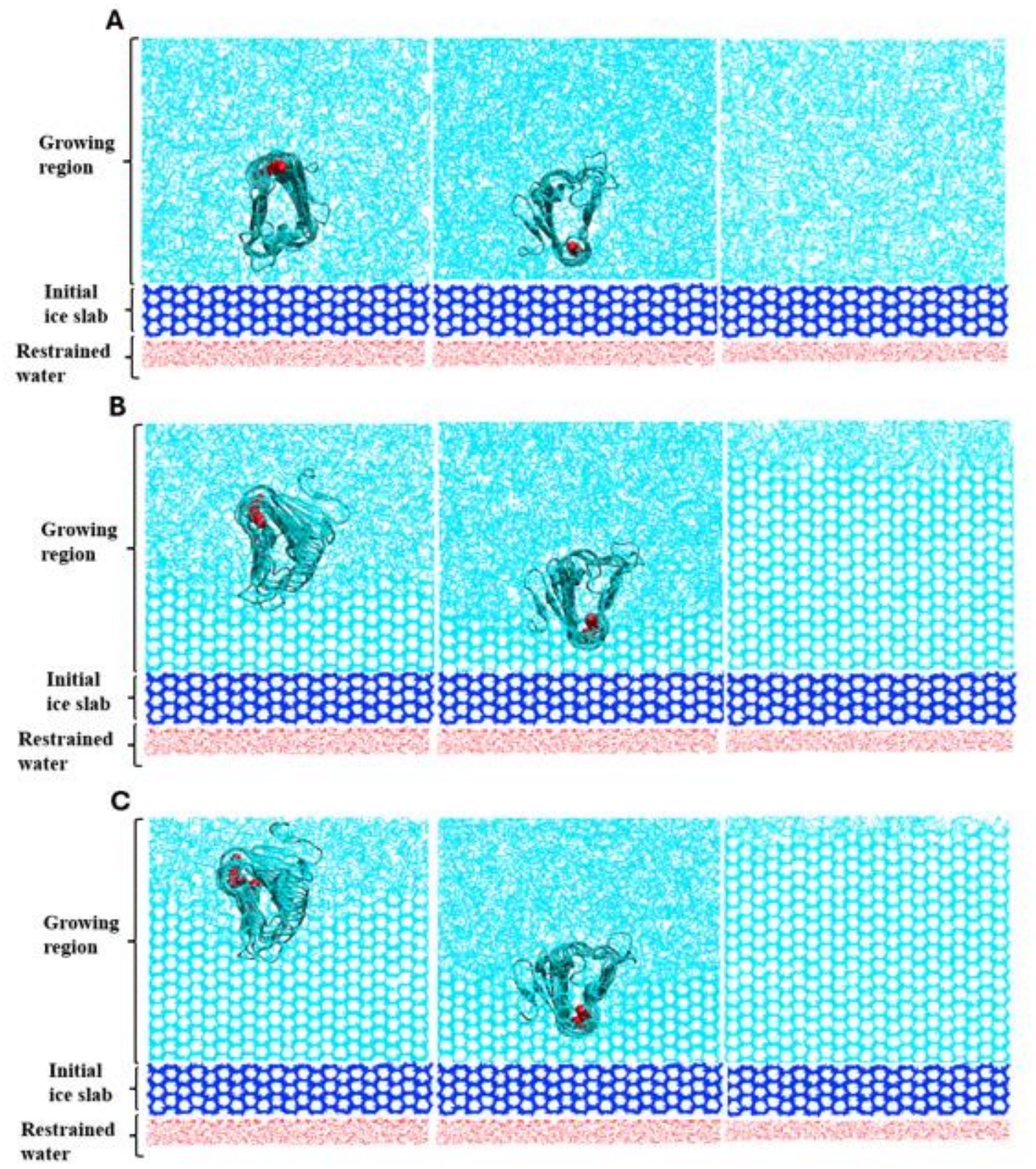
Representative simulation trajectories of ice growth on the primary prism plane for the non-IBS-exposed MaIBP_RIV, IBS-exposed MaIBP_RIV AFP, and ice/water system without any protein (top to bottom in the chronological order) at (A) 0 ns, (B) 100 ns, and (C) 1000 ns (for the system without the protein, it was ~200 ns) were shown. The preformed ice slab was rendered in blue, and the oxygen atoms of water molecules were shown in cyan colour. The position-restrained water molecules were shown in red colour. Protein was rendered in new cartoon mode. Ca^2+^ ions were shown in VDW mode and in red colour.

When the IBS of MaIBP_RIV was exposed, the growing ice front formed a stable contact with the IBS. Once adsorption occurred, the protein strongly bound to the ice surface throughout the simulation (Figure 2). This adsorption effectively inhibited further progression of the ice front. This observation was consistent with the adsorption-inhibition mechanism, in which AFPs bind to the ice surface and create a curved ice-water interface between adjacent bound proteins (Figure S3A, generated using VMD, with periodic image applied). When the radius of this curved interface becomes smaller than the critical radius required for ice formation, further ice growth is inhibited. No further progression of the ice front was observed after binding. This indicated a complete suppression of ice growth. Notably, this arrested growth state persisted for the entire 1000 ns simulation (Figure 2), suggesting a strong affinity of the IBS for the primary prism plane and supporting the role of adsorption-mediated inhibition in the antifreeze activity of MaIBP_RIV.

In comparison, when the non-IBS side of MaIBP_RIV was exposed, it exhibited different behaviour. Rather than forming stable interactions with the ice surface, the protein was continuously repelled by the growing ice front (Figure 2). The presence of the protein decreased the overall rate of ice growth compared to the pure ice/water system. However, it was unable to halt the formation of ice. Furthermore, it also showed continuous rotational motion during the simulations (Figure 2). This suggests the absence of a specific ice-binding surface to form stable interactions with the growing ice surface. These observations indicated that the inhibition of ice growth by MaIBP_RIV was due to specific molecular recognition mediated by the IBS.

To quantify ice growth, the fraction of ice-like water molecules was calculated using the CHILL+ order parameter. In all systems, the fraction of ice-like molecules increased with simulation time. However, the extent of growth differed significantly among the systems (Figure 5A). The ice/water system reached nearly 0.8 fraction of ice-like molecules within 100 ns, indicating unrestricted growth (Figure 5A). But the fraction of ice-like molecules for the non-IBS-exposed case reached approximately 0.5 within 500 ns (Figure 5A) and plateaued after that. Careful inspection reveal that this plateauing is not due to stoppage of ice growth but because of the presence of protein in the system. The non-IBS side is repelled by the advancing ice front and does not establish stable binding with the growing ice surface. Instead, the plateau is mainly associated with the presence of the protein near the end of the simulation box. The protein occupies a certain volume and limits the space available for further ice formation. Therefore, the non-IBS reduced ice growth to some extent but failed to prevent continuous formation of ice (Figure 5A). The fraction of ice-like molecules continued to increase towards the highest possible limit, indicating continuous ice growth. In contrast, the IBS-exposed system reached a fraction of ice-like molecules of nearly 0.2 within 500 ns (Figure 5A). After adsorption of the AFP to the ice surface, the growth curve reached a plateau till 1000 ns, indicating suppression of further ice growth (Figure 5A). The plateau provides the evidence for the strong binding affinity of MaIBP_RIV toward the primary prism plane and its ability to effectively arrest ice growth. Two independent sets of simulations were performed for each system, including the pure ice/water, non-IBS-exposed, and IBS-exposed MaIBP_RIV. The results from another set were provided in the Supporting Information (Figure S2A). Importantly, the similar behaviour was observed across two simulation sets.

### 3.3. Ice-Growth Inhibition on the Basal Plane

To examine whether the inhibition behaviour of MaIBP_RIV was maintained on other faces of ice, independent simulations were performed at the basal plane of hexagonal ice. Comparative simulation snapshots for the IBS-exposed MaIBP_RIV, non-IBS-exposed MaIBP_RIV, and pure ice/water systems were presented in Figure 3. Similar to the observations on the primary prism plane, the IBS of MaIBP_RIV was exhibited strong binding to the growing ice front. During this adsorption, the protein remained stably associated with the ice surface and effectively inhibited further ice growth on the basal plane. This adsorption was consistent with the adsorption-inhibition mechanism, in which AFPs bind to the ice surface and create a curved ice-water interface between adjacent bound proteins (Figure S3B, generated using VMD, with applying periodic image). As the radius of this curved water interface becomes smaller than the critical radius required for ice formation, further ice growth is inhibited. The suppression of further ice growth throughout the simulation indicated the strong affinity of IBS of MaIBP_RIV for the basal plane.

**Figure 3:**
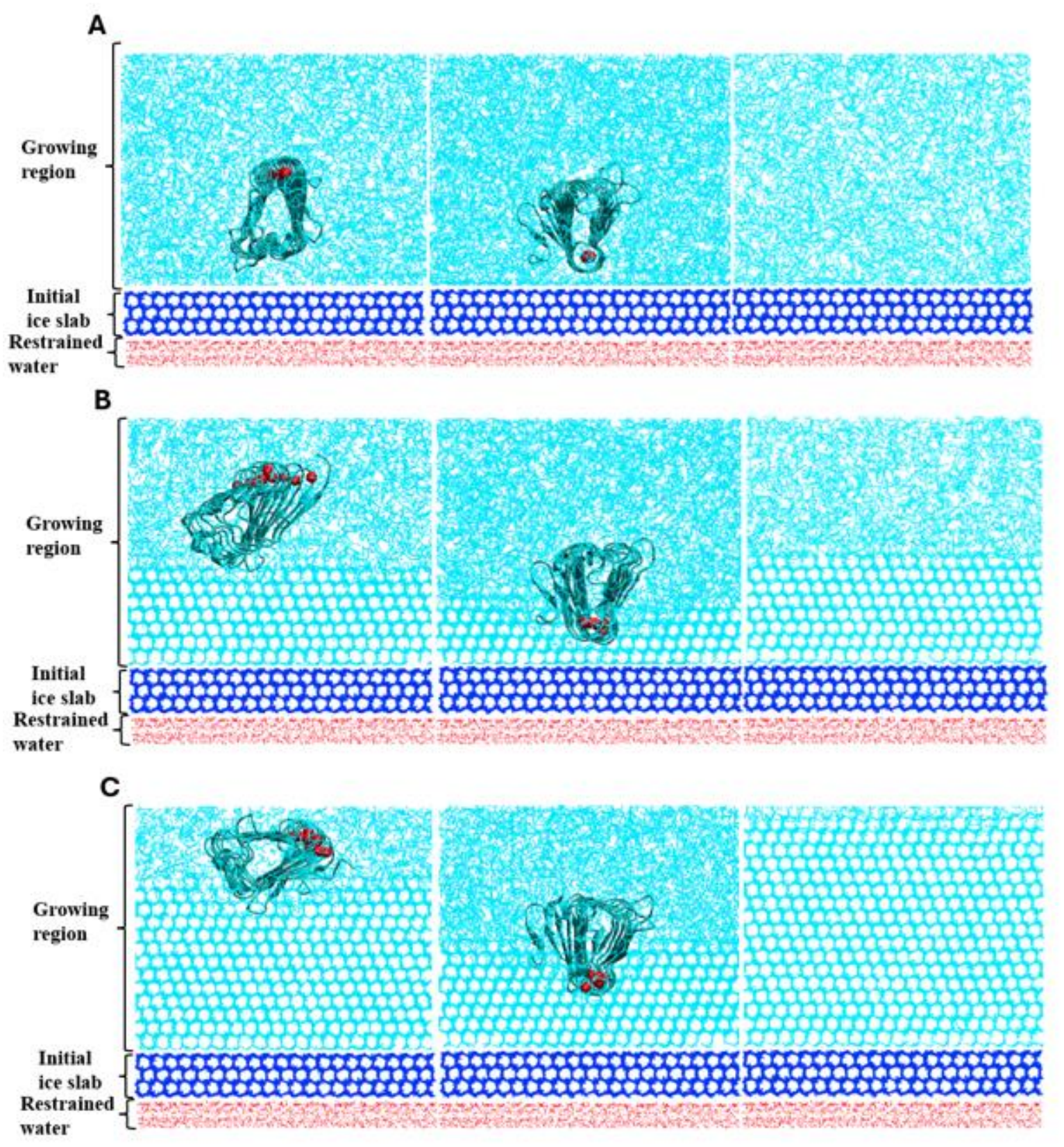
Representative structures from the simulation trajectories of ice growth on the basal plane for the non-IBS-exposed MaIBP_RIV, IBS-exposed MaIBP_RIV, and the system without any protein (top to bottom in chronological order) were shown. Ice growth occurs along the basal plane. Snapshots of representative images for all three systems obtained from the production molecular dynamics simulation at (A) 0 ns, (B) 100ns, and (C) 1000 ns (for the system without protein, it was 250 ns) were shown. Preformed ice was represented in blue, and oxygen atoms of the water molecules were shown in cyan colour. The position-restrained water molecules were shown in red colour. Protein was represented in new cartoon mode. Ca^2+^ ions were shown in VDW mode and in red colour.

The behaviour of the non-IBS of MaIBP_RIV on the basal plane was closely similar to that observed on the primary prism plane. In Figure 3, the growing ice front continuously repelled the protein to prevent the formation of stable protein-ice interactions. Therefore, the non-IBS was unable to arrest ice growth (Figure 3). It also showed substantial rotational and translational motion throughout the simulation (Figure 3). However, a modest reduction in ice growth was observed relative to the pure ice/water system. So, the non-IBS did not exhibit the strong growth inhibition properties like the IBS of MaIBP_RIV (Figure 3).

Quantitative analysis using the CHILL+ order parameter further confirmed these observations (Figure 5B). In Figure 5B, the fraction of ice-like molecules reached nearly 0.7 within 220 ns in the ice/water system, reflecting unrestricted ice growth. The non-IBS-exposed system produced only partial inhibition, resulting in a continuous increase in the ice-like water population to approximately 0.5 (Figure 5B) and plateaued after that. A closer look shows that the plateau does not occur because ice growth has inhibited. Like the primary prism plane, the plateau mainly occurs because the protein is located near the end of the simulation box. Thus, the non-IBS slowed down the ice growth to some extent but could not stop it completely (Figure 5B). In comparison, the IBS-exposed system substantially reduced the formation of ice-like molecules and plateaued at a value of nearly 0.2 in the simulation (Figure 5B). The growth curve approached a plateau after binding of the protein to the ice surface (Figure 5B). These results demonstrated that stable binding of the IBS of MaIBP to the basal plane effectively suppresses ice formation. For each system, including pure ice/water, non-IBS-exposed, and IBS-exposed MaIBP_RIV, two independent simulations were performed. The results of another set are provided in the Supporting Information (Figure S2B). Similar phenomena were observed in both sets.

### 3.3. Ice-Growth Inhibition on the Secondary Prism Plane

To examine whether the ice-growth inhibition by MaIBP_RIV was maintained on the secondary prism plane, simulations were performed at the secondary prism plane of hexagonal ice. Representative snapshots of the non-IBS-exposed, IBS-exposed and pure ice/water systems were shown in Figure 4. Here, IBS-exposed MaIBP_RIV showed weaker binding to the growing ice front on the secondary prism plane compared with the primary prism and basal planes. Interestingly, MaIBP_RIV effectively reduced ice growth rate during the initial and intermediate stages of the simulation. The adsorption was consistent with the adsorption-inhibition mechanism, in which AFPs bind to the ice surface and create a curved ice-water interface between adjacent bound proteins (Figure S3C, generated using VMD, with periodic image applied). The formation of this curved interface initially restricted the progression of the ice front. At later stages, ice growth gradually resumed, as indicated by the increase in the fraction of ice-like molecules, suggesting that the weaker binding to the secondary prism plane was insufficient to maintain complete inhibition of ice growth throughout the simulation.

**Figure 4:**
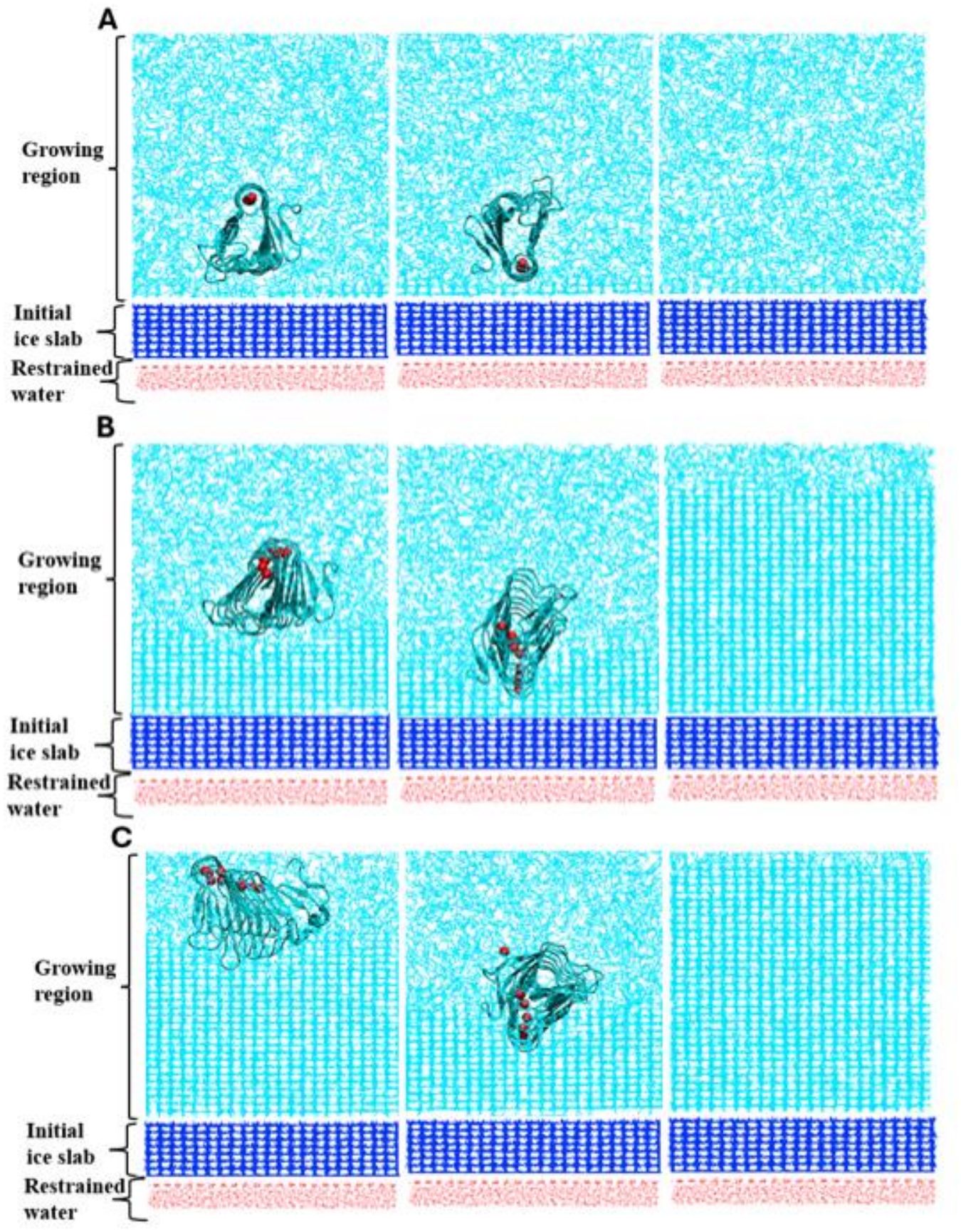
Representative simulation trajectories of ice growth on the secondary prism plane for the non-IBS-exposed MaIBP_RIV, IBS-exposed of MaIBP_RIV, and ice/water system without any protein (top to down in the chronological order) at (A) 0 ns, (B) 100 ns, and (C) 1000 ns (for the system without the protein, it was ~150 ns) were shown. The preformed ice slab was rendered in blue, and the oxygen atoms of water molecules were shown as cyan colour. The position restrained water molecules were shown in red colour. Protein was rendered as a new cartoon mode. Ca^2+^ ions were shown in VDW mode and in red colour.

The non-IBS-exposed system showed behaviour similar to that observed on the primary prism and basal planes. During the simulation, the growing ice front continuously repelled the protein. As a result, the non-IBS was unable to arrest ice growth and showed considerable rotational and translational motion throughout the simulation. Although the protein slightly reduced the rate of ice growth compared with the pure ice/water system, it did not show strong ice-growth inhibition (Figure 4).

Quantitative analysis using the CHILL+ order parameter further confirmed these observations (Figure 5C). In Figure 5C, the fraction of ice-like molecules reached nearly 0.85 within 200 ns in the ice/water system, reflecting unrestricted growth. The non-IBS-exposed system produced only partial inhibition, resulting in a continuous increase in the fraction of ice-like molecules to approximately 0.5 (Figure 5C) and plateaued after that. The non-IBS side was pushed away by the advancing ice front. Similar to the primary prism and basal plane, the plateau appears to result from the protein’s presence at the end of the simulation box. As a result, it slowed down ice growth to some extend but could not stop it completely (Figure 5C). In comparison, the IBS-exposed system substantially reduced the formation of ice-like molecules. The fraction initially remained near 0.1 before gradually increasing to approximately 0.4 toward the end of the simulation (Figure 5C). The growth curve showed a slow increase in the fraction of ice-like molecules throughout the simulation (Figure 5C). Therefore, the IBS of MaIBP_RIV reduces ice growth on the secondary prism plane, but it did not completely supress ice formation. However, the suppression is relatively better compared to non-IBS-exposed system. We also observed a similar scenario in another set of simulation and presented the results in Supporting Information (Figure S2C).

**Figure 5:**
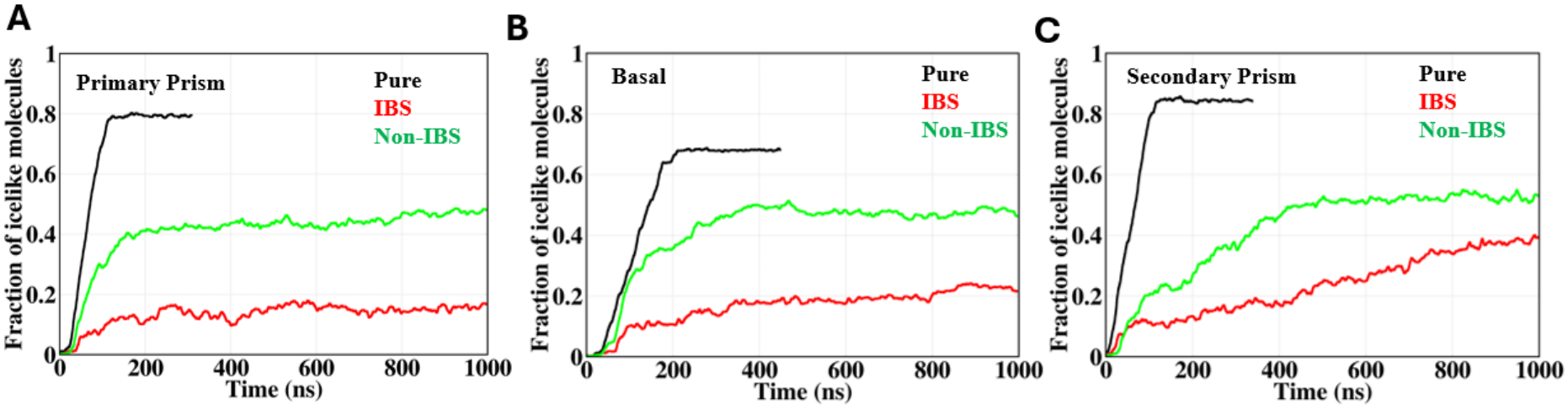
The evolution of ice like ordered water in the ice growth simulation of non-IBS-exposed MaIBP_RIV, IBS-exposed MaIBP_RIV and the pure ice/water system with time on three different ice surfaces (A) primary prism plane, (B) basal plane, and (C) secondary prism plane.

### 3.4. The Importance of Ca^2+^ Ion for IBS Stability of MaIBP_RIV AFP

Next, we examine the role of Ca^2+^ in structure and function of MaIBP_RIV. In Figure 6A, the results clearly depicted the significant role of Ca^2+^ ions in maintaining the ice-binding surface (IBS) stability of MaIBP_RIV and enhanced its antifreeze activity. Two independent simulation sets were performed for each system. Here, we presented the results from one representative simulation set, while the results from another set were provided in the Supporting Information (Figure S4). In the absence of any protein, the ice/water system exhibited rapid ice growth. The fraction of ice-like molecules increased linearly during the first 100 ns on the primary prism plane. It reached a plateau of approximately 0.8 by around 120 ns (Figure 6A). In comparison, when the IBS of MaIBP_RIV AFP with Ca^2+^ was exposed, it exhibited the strongest suppression of ice growth mechanism. Throughout the simulation, the fraction of ice-like molecules remained considerably lower than in the pure ice/water system. The fraction of ice-like molecules reached only about 0.2 (Figure 6A). But, when IBS of MaIBP_RIV without Ca^2+^ ions was exposed to ice, the ice growth rate was reduced compared with the pure ice/water system on the primary prism plane. However, the inhibition remained relatively weak. The fraction of ice-like molecules gradually increased to approximately 0.5 after 400 ns and remained the same throughout the simulation. Careful inspection reveal that this plateauing is not due to stoppage of ice growth but because of the presence of protein in the system. The observed plateau is likely due to the presence of the protein near the end of the simulation box. Since the protein occupies some space, it reduces the area available for further ice formation. Therefore, the IBS of MaIBP_RIV without Ca^2+^ exposed slowed down ice growth to some extent, it could not completely stop ice formation (Figure 6A). The fraction of ice-like molecules continued to increase and approached its maximum value, indicating that ice growth continued throughout the process. This value was nearly half that observed for the ice/water system. The same overall behaviour was observed in other sets also (Figure S4A).

**Figure 6:**
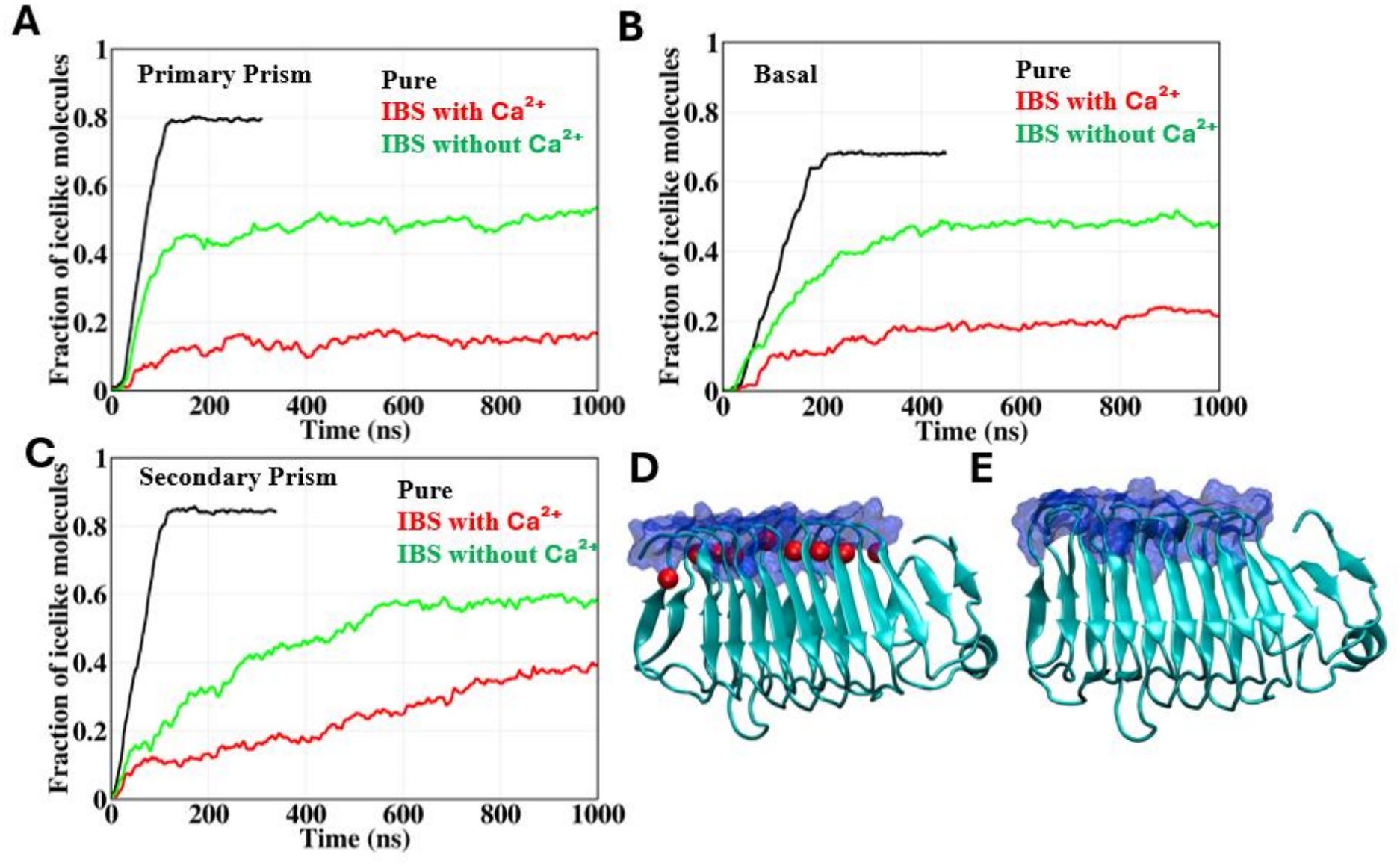
Time evolution of the fraction of ice-like molecules on the primary prism plane (A), basal plane (B), and secondary prism plane (C) respectively for three systems: ice/water without protein (black), Ca^2+^ bound MaIBP_RIV (red), and MaIBP_RIV without Ca^2+^ bound (green). The structure of Ca^2+^ bound MaIBP_RIV in the IBS (D) & the structure of MaIBP without Ca^2+^ bound in the IBS (E). The IBS residues were shown in the surface mode in blue colour (D and E) and Ca^2+^ ions were shown in VDW mode and in red colour.

In the pure ice/water system, ice growth formation occurred continuously throughout the simulation on the basal plane (Figure 6B). It reached a plateau of about 0.68 at around 220 ns (Figure 6B). The fraction of ice-like molecules indicated extensive growth on the basal plane in the absence of any protein. In comparison, when the IBS of MaIBP with Ca^2+^ was exposed, it exhibited the strong inhibition of ice growth on the basal plane. The fraction of ice-like molecules remained very low throughout the simulation and reached only approximately 0.20 (Figure 6B). However, when the IBS of MaIBP_RIV without Ca^2+^ was exposed, the ice-growth inhibition ability was reduced compared to pure ice/water system. The fraction of ice-like molecules gradually increased to approximately 0.5 and plateaued after that (Figure 6B). A closer look shows that the plateau does not occur because ice growth has inhibited. The plateau is mainly due to the presence of the protein at the far end of the simulation box. Like the primary prism plane the IBS of MaIBP_RIV without Ca^2+^ exposed partially suppressed ice growth but was unable to completely inhibit the continuous formation of ice (Figure 6B). Similar behaviour was observed in the other simulation (Figure S4B)

On the other hand, the pure ice/water system, ice growth continued throughout the simulation on the secondary prism plane. The fraction of ice-like molecules increased steadily after an initial period of approximately 30-40 ns. The value of fraction reached approximately 0.85 at around 200 ns (Figure 6C). This high fraction of ice-like molecules indicated extensive ice growth in the absence of protein. In contrast, IBS of MaIBP_RIV with Ca^2+^ exposed showed inhibition of ice growth on the secondary prism plane. The fraction of ice-like molecules remained relatively low during the initial stage of the simulation, at approximately 0.1. But, the value of fraction of ice-like molecules gradually increased at later stages and reached approximately 0.4 by the end of the simulation (Figure 6C). In comparison, when the IBS of MaIBP_RIV without Ca^2+^ exposed, it showed a weaker ability to inhibit ice growth. The fraction of ice-like molecules continued to increase throughout the simulation and reached approximately 0.6 by 700 ns (Figure 6C) and remained the same throughout the simulation. As observed for the primary prism and basal planes, the plateau occurred because the protein was located near the end of the simulation box. Therefore, the IBS of MaIBP_RIV without Ca^2+^ exposed partially slowed down ice growth, it was unable to completely prevent further ice formation (Figure 6C). However, this value was lower than that of the pure ice/water system. The continuous increase of the ice-like molecules indicated that Ca^2+^ free MaIBP_RIV could not effectively stop ice growth. Compared with both the pure ice/water and Ca^2+^ free MaIBP_RIV systems, the lower fraction of ice-like molecules indicated that Ca^2+^ bound MaIBP_RIV reduced ice growth in a better way. However, unlike the primary prism and basal plane, ice growth was not completely stopped during the simulation. These results suggested that Ca^2+^ helped to maintain the structural stability of MaIBP_RIV and enhanced its ice-binding ability. The same overall behaviour was observed in other sets also (Figure S4C). However, the extend of inhibition of the ice growth process depends on the ice surface.

In Figure 6D, the structure of MaIBP_RIV with Ca^2+^ was shown during the simulation at 315 ns on the primary prism plane. It highlighted the flatness of the IBS. The IBS remained flat and remained intact throughout the simulation in the presence of the Ca^2+^ ions. In the presence of Ca^2+^, the RMSD of IBS residues remained approximately 0.15 nm (Figure S5). The structure of MaIBP_RIV without Ca^2+^ from simulation at time 98 ns on the primary prism plane was presented in Figure 6E. The flatness of the IBS of MaIBP_RIV was deformed in the absence of Ca^2+^ ions. In the absence of Ca^2+^, the RMSD of IBS residues increased to ~0.25 nm (Figure S5). Here, we concluded that Ca^2+^ ions are important for maintaining flatness of the IBS of MaIBP_RIV which in tun is important for proper binding with the different plane of hexagonal ice.

### 3.5. Effect of Ice Binding Surface (IBS) Mutations on Ice Growth Inhibition

To further investigate the effect of mutations on the IBS of MaIBP_RIV on ice-growth inhibition, a series of site-directed mutations were introduced. Four mutants, S44Y, G63D, N65Y, and D87Y were selected based on the experimental results. The experimental results showed a reduction in antifreeze activity due to these mutations. The simulations were performed on the primary prism plane, as it demonstrated strong and stable binding to this plane. The ability of each mutant to inhibit ice growth was assessed by monitoring the fraction of ice-like molecules as a function of simulation time.

Figure 7A compared the evolution of the fraction of ice-like molecules for the wild-type MaIBP_RIV and four IBS mutants: S44Y, N65Y, D87Y, and G63D. The wild-type protein exhibited the strongest ice-growth inhibition among all systems. After an initial increase during the first 100 ns, the fraction of ice-like molecules stabilized at approximately 0.20 and remained relatively constant throughout the simulation (Figure 7A). This behaviour indicated strong binding of MaIBP_RIV onto the ice surface and inhibited the ice-growth process.

**Figure 7:**
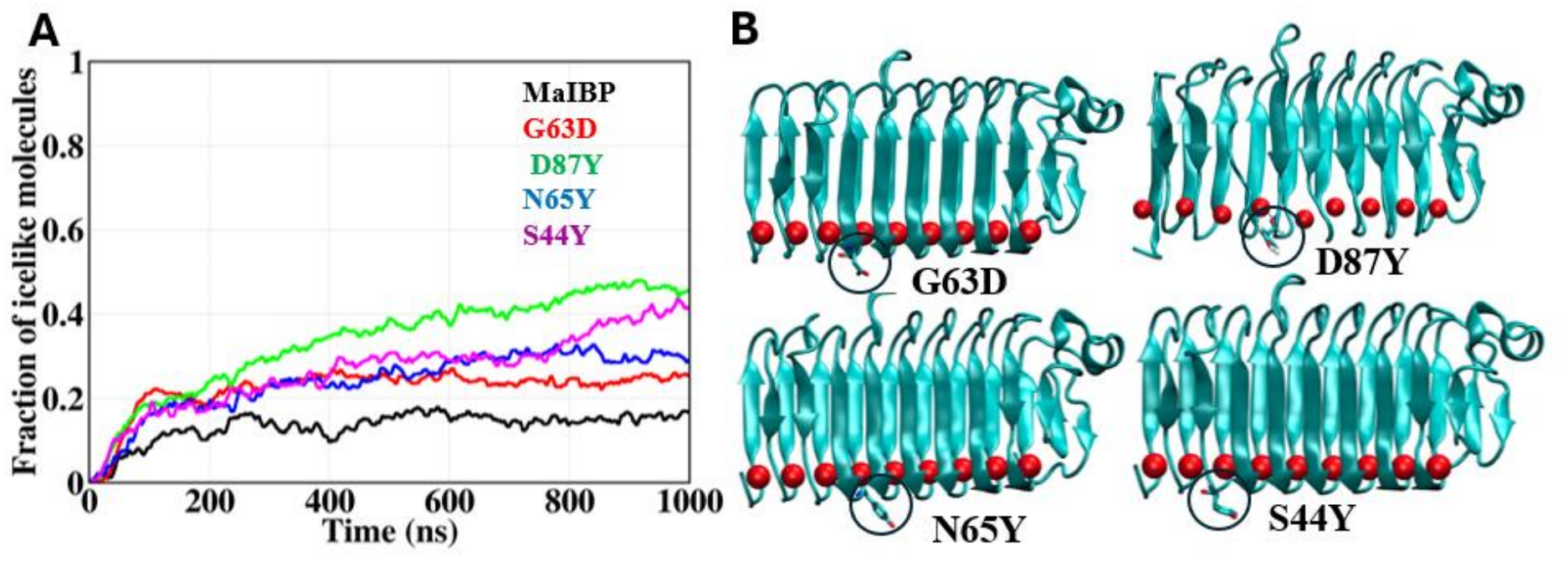
Time evolution of the fraction of ice-like molecules on the primary prism plane for wild-type MaIBP_RIV (A) and four ice binding surface mutants (G63D, D87Y, N65Y, and S44Y) (B). Ca^2+^ ions were shown in VDW mode and in red colour.

In contrast, all IBS mutants showed a higher fraction of ice-like molecules than the wild-type protein and more importantly they showed slow but somewhat continuous growth. This indicated a reduction in antifreeze activity upon mutation of key IBS residues. This result explained that the native arrangement of residues within the IBS was crucial for maintaining efficient ice-growth inhibition. Among the mutants, D87Y exhibited the most pronounced loss of function. The fraction of ice-like molecules continuously increased throughout the simulation and reached approximately 0.4 by the end of the trajectory (Figure 7A). The absence of a plateau suggested persistent ice-growth. It indicated that Asp87 played a crucial role in maintaining the structural and functional integrity of the ice-binding interface. Substitution of this residue with tyrosine likely disrupted interactions between the IBS and the ice surface and significantly weakened the binding on the ice surface. The S44Y mutant also showed a substantial reduction in ice-growth inhibition. The ice fraction continued to increase steadily and reached approximately 0.4 (Figure 7A). The gradual increase suggested that mutation of Ser44 compromised the stability of the protein-ice interaction. So, it allowed the ice-growth process on the primary prism plane. Next, the N65Y and G63D mutants also showed a gradual increase in the fraction of the ice-like molecules. The value of the fraction of ice-like molecules reached approximately 0.3 during the simulation (Figure 7A). Overall, a continuous increase in the ice-like fraction was observed for all four mutants. This indicates that these IBS residues contribute to maintaining effective ice-growth inhibition by MaIBP_RIV.

### 3.6. Hydration Structure Around the Ice-Binding and Non-Ice-Binding Surface of MaIBP_RIV

To characterize the structural organization of water molecules surrounding the ice-binding surface (IBS) of MaIBP_RIV antifreeze protein (AFP), the F_4_ was calculated over the temperature range of 240-300 K. For each temperature, 200 ns molecular dynamics simulations were performed. The first 20 ns were excluded for the calculation. All hydration analyses were calculated using the final 180 ns of trajectories. Negative F_4_ values correspond to ice-like water ordering, whereas positive F_4_ values indicate the formation of more clathrate-like hydration structures.

Figure 8 illustrates the variation of F_4_ around IBS and non-IBS of MaIBP_RIV from 300 K to 240 K. At 300 K, both IBS and non-IBS exhibited F_4_ values close to zero, with IBS and non-IBS having values of approximately −0.010 and −0.005, respectively. These values indicated that the water molecules around IBS and non-IBS were liquid like. As the temperature decreased, the F_4_ values became more negative. The decrease of F_4_ values were more pronounced for the IBS of MaIBP_RIV, where F_4_ decreased from approximately −0.010 at 300 K to −0.055 at 240 K. But, the non-IBS shows a comparatively moderate decrease from approximately −0.005 to −0.022 over the temperature range from 300K to 240K. This trend indicated that decreasing the temperature enhanced the ordering of water molecules particularly around the IBS of MaIBP_RIV. Throughout the temperature range, the IBS consistently exhibited more negative F_4_ values than the non-IBS. These explained that the water molecules around the IBS maintained an ice-like hydration environment with decreasing temperature from 300K to 240K.

**Figure 8:**
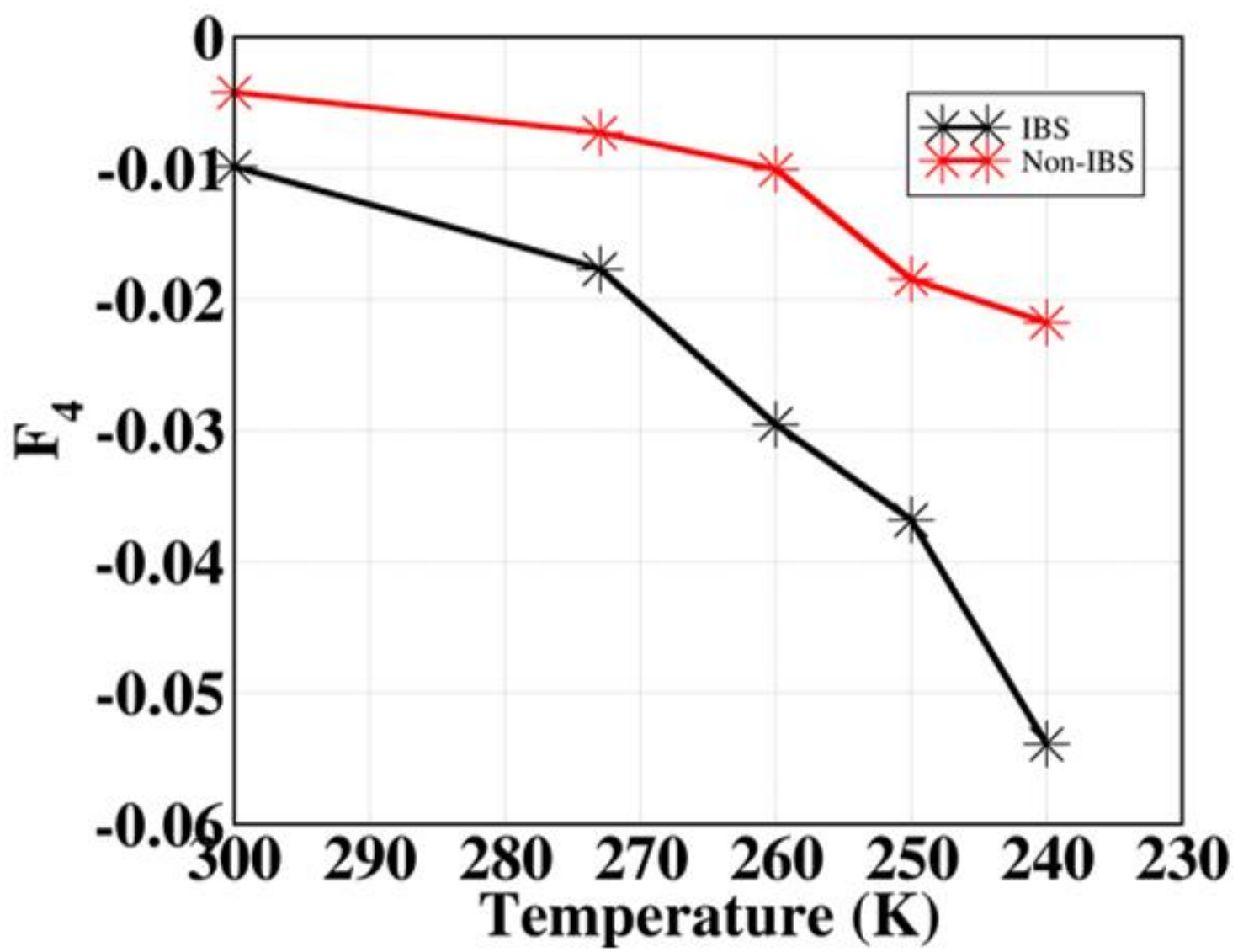
Temperature dependence of F_4_ for the ice-binding surface (IBS) and non-ice-binding surface (non-IBS) of MaIBP_RIV.

## 4. CONCLUSION

The present molecular dynamics study provided detailed mechanistic insights into the antifreeze activity of MaIBP_RIV. This study revealed the molecular details regarding its ice growth inhibition properties. Simulations performed at both the primary prism and basal planes of hexagonal ice. The results explained that MaIBP_RIV effectively suppressed ice-growth through specific binding of the ice-binding surface (IBS) onto the growing ice surface. The protein remained strongly bound to the ice surface and completely arrested further ice-growth throughout the simulation timescale. On the other hand, the non-IBS side of MaIBP_RIV failed to establish stable interactions with growing ice and exhibited substantial rotational and translational motion. Therefore, non-IBS was unable to prevent the ice-growth process. The simulations additionally revealed critical role of Ca^2+^ ions in maintaining antifreeze activity. Removal of Ca^2+^ significantly weakened the ability of MaIBP_RIV to inhibit ice-growth. Ca^2+^ was found to be essential for preserving the flat IBS structural topology important for establishing stronger binding to the ice surface. Different site-directed mutagenesis studies further identified key residues involved in the growth inhibition. Hydration analyses based on the F_4_ order parameter revealed that the wild-type IBS promoted more ice-like ordering at lower temperature than non-IBS.

Overall, this study demonstrated that the antifreeze activity of MaIBP_RIV was governed by the specific IBS residues, Ca^2+^ dependent structural stabilization, and the formation of an ordered hydration structure at the protein surface. Stable binding of the protein onto ice effectively arrested the ice-growth and prevented formation of the ice phase. These findings provide a comprehensive molecular understanding of MaIBP_RIV function and offer valuable insights for the rational design of bioinspired antifreeze materials and cryoprotective agents.

### Notes

The authors declare no competing financial interest.

## ACKNOWLEDGEMENTS

This research was supported by the Department of Science and Technology ANRF grant ANRF/ARG/2025/003888/CS for funding. SS thanks IACS for the fellowship and computational facility.

