## Supplementary Information for "Insights into Ice-Growth Inhibition and Hydration Ordering by the Ca^2+^-Dependent Antifreeze Protein MaIBP_RIV"

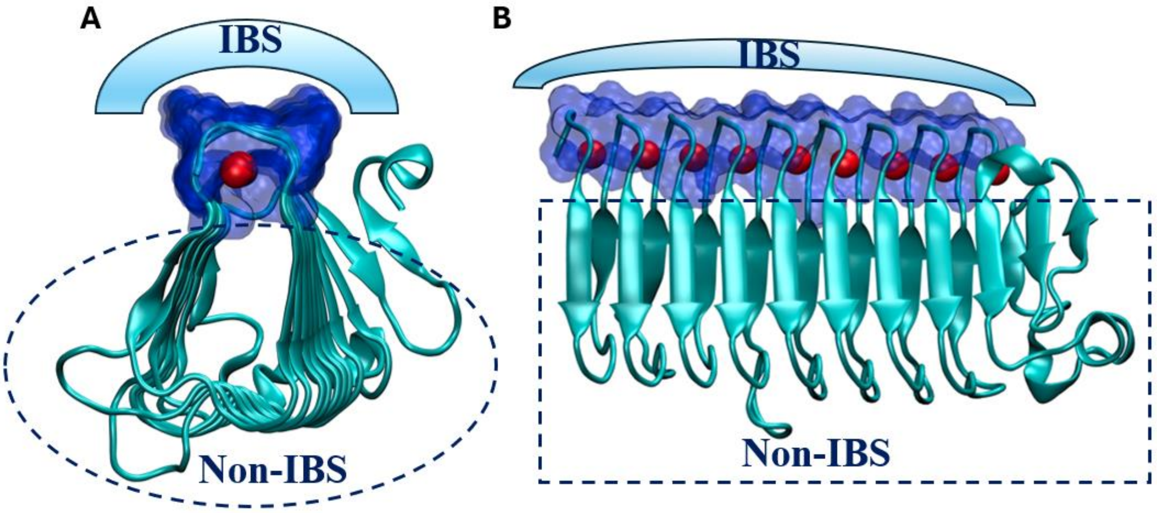


S1: Structural representation of the ice-binding surface (IBS) of MaIBP_RIV. (A) Front view of the MaIBP_RIV and (B) Side view of the MaIBP_RIV. The protein structure was shown in new cartoon representation, IBS in surface mode (indicated by the shaded region) and $Ca^{2+}$ ions in VDW (red colour) respectively.

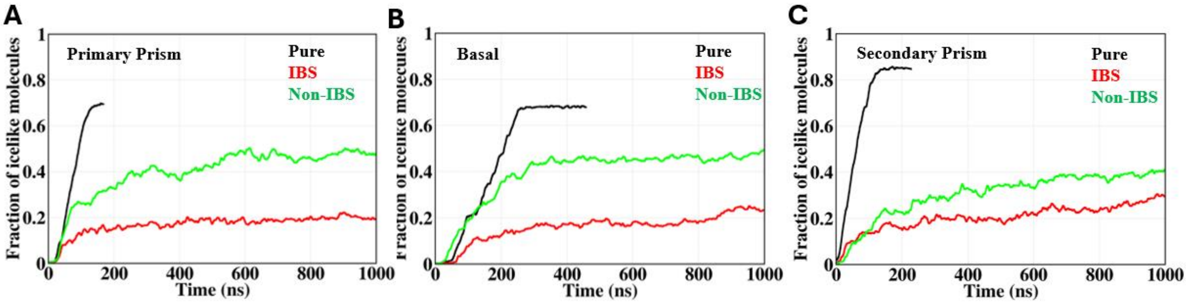

S2: The evolution of ice like ordered water in the ice growth simulation of non-IBS exposed MaIBP_RIV, IBS exposed MaIBP_RIV and the pure ice/water system with time on three different ice surfaces (A) primary prism plane, (B) basal plane, and (C) secondary prism plane.

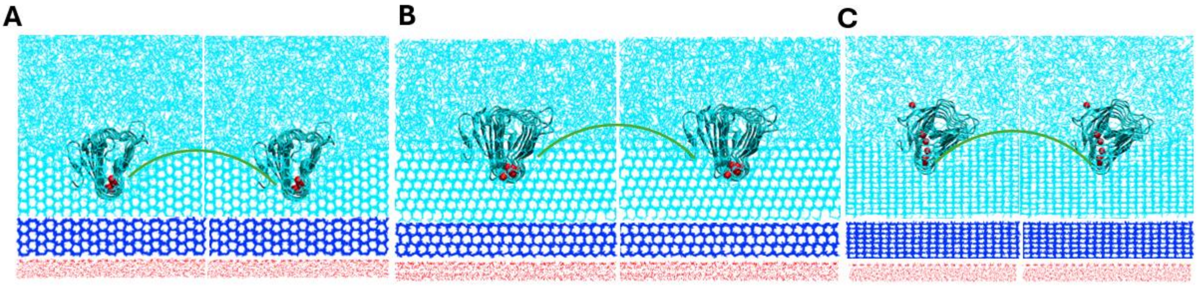


S3: Formation of a curved surface after the binding of MaIBP_RIV on the primary prism plane (A), basal plane (B) and secondary prism plane (C) observed during the ice growth simulation. Oxygen atoms of the water molecules were represented in cyan colour. The preformed ice slab was rendered in blue, and the oxygen atoms of water molecules were shown as cyan colour. The position restrained water molecules were shown in red colour. Protein was rendered as a new cartoon mode. $Ca^{2+}$ ions were shown in VDW mode and in red colour.

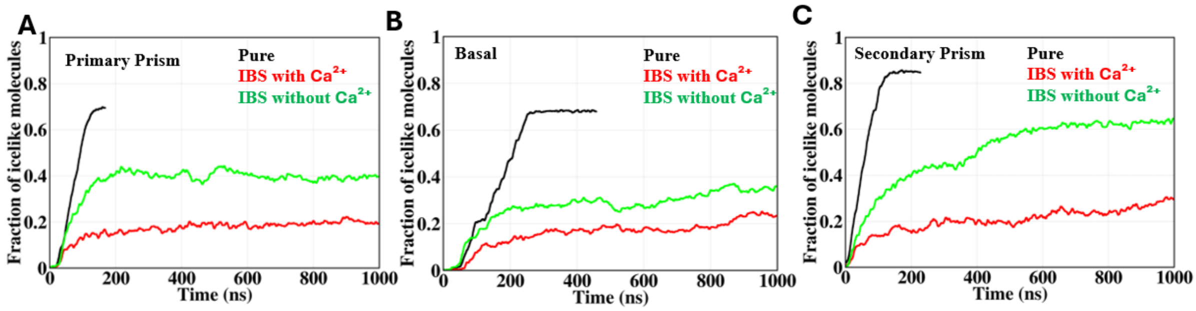

S4: Time evolution of the fraction of ice-like molecules on the primary prism plane (A), basal plane (B), and secondary prism plane (C) respectively for three systems: ice/water without protein (black), $Ca^{2+}$ bound MaIBP_RIV (red), and MaIBP_RIV without $Ca^{2+}$ bound (green).

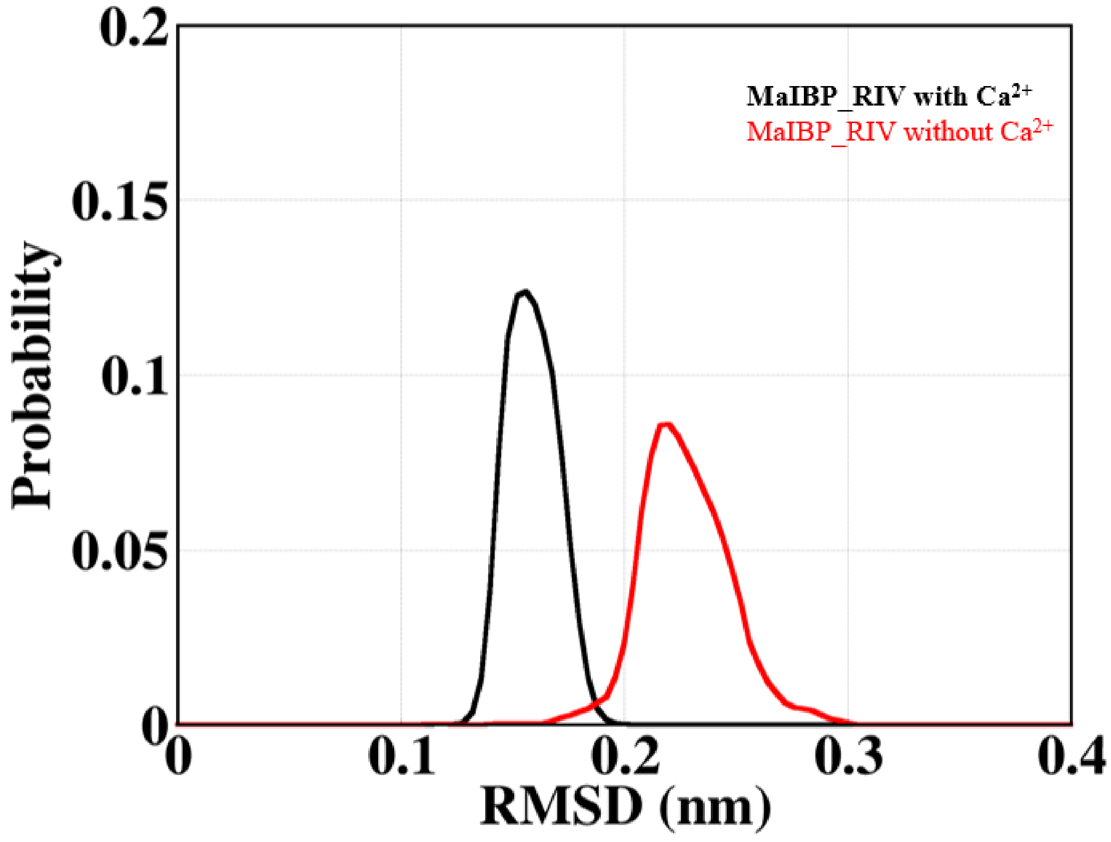


S5: RMSD distribution of the IBS residues of MaIBP_RIV in the presence (black) and absence (red) of $Ca^{2+}$.
